# Mice developing mammary tumors evolve T cell sequences shared with human breast cancer patients

**DOI:** 10.1101/371260

**Authors:** Miri Gordin, Hagit Philip, Alona Zilberberg, Moriah Gidoni, Raanan Margalit, Christopher Clouser, Kristofor Adams, Francois Vigneault, Irun R. Cohen, Gur Yaari, Sol Efroni

## Abstract

Cancer immunotherapy by checkpoint blockade proves that an effective immune response to a tumor can be induced clinically. However, little is known about the evolution of tumor-associated T-cell receptor (TCR) repertoires without intervention. Here we studied TCR repertoire evolution in mice spontaneously developing mammary tumors; we sequenced peripheral blood alpha and beta TCRs of CD4^+^CD62L^+^CD44^−^ T cells monthly for 8 months in 10 FVB/NJ mice transgenic at the Erbb2 locus, all developing tumors; 5 FVB/NJ mice without the transgene were age-matched controls. Sequences were either private (restricted to one mouse) or public (shared among mice); public sequences were either exclusive to the tumor group or inclusive among different groups. We now report that 1), public AA sequences were each encoded by many different nucleotide sequences (NT) recombinations (convergent recombination; CR); 2) mice developing tumors evolved tumor-exclusive public sequences, derived initially from private or from inclusive public sequences; and 3) tumor-exclusive public sequences in mice were also present among published public TCR sequences from human breast cancer patients. These cross-species tumor-exclusive TCR sequences manifested high CR; but the AA sequences shared by mice and humans did not share NT sequences. Thus, tumor-exclusive TCR AA sequences across species are selected from different NT recombination events. The roles of tumor-exclusive TCR repertoires in advancing or inhibiting tumor development and the effects of tumor immunotherapy on these T cells remain to be seen.

## Introduction

T cells control multiple immune mechanisms that are involved in tumor progression (Dunn, Bruce, Ikeda, Old, & Schreiber, 2002; Hanahan & Weinberg, 2011). Attempts to influence the interaction between T cells and cancer cells often rely on specific peptide presentation on the surface of tumor cells or of antigen presenting cells (Butterfield, 2015; Kawakami et al., 1994; Simpson, Caballero, Jungbluth, Chen, & Old, 2005; Slingluff, 2011), the presentation of a relevant receptor on the surface of an interacting T cell (Brentjens et al., 2013; Ye et al., 2018), or on the manipulation of co-stimulation of the interaction (Drake, Lipson, & Brahmer, 2014; Hoos et al., 2010; Pardoll, 2012; Sonpavde, 2017). Each T cell clone is characterized by a heterodimeric T-cell Receptor (TCR), and its interaction with a (tumor) cell is further characterized by a protein degradation product presented by the major histocompatibility complex (MHC) on the cell’s surface. For most T-cells, the heterodimeric TCR is composed of α (TRA) and β (TRB) chains (less than 10% express the γ and δ form of TCR(Kalyan & Kabelitz, 2013)). The sequences of TRB chains are ultimately determined by somatic V(D)J recombination of the Variable (V), Diversity (D) and Joining (J) paralogs, and the recombination of VJ for the TRA chain. This recombination includes addition, deletion, and replacement of nucleotides, which leads to extreme possible variability of the complementarity-determining region 3 (CDR3) loop, that interacts with the MHC presented peptides (Pellicci et al., 2014). The collection of available receptors on the surface of T cells has been termed the T-cell repertoire (Fukui et al., 1998). T cells make up to 60% of PBMCs, so that even small volumes of drought peripheral blood, contain a large number of T cells, such that a milliliter of peripheral blood contains 0.5-1.5 million T cells (Pagana & Pagana, 2007). T cells are early responders to neoplasm, and we set out to find if changes to the T cell repertoire, previously demonstrated in tumor (Galon et al., 2006; Ino et al., 2013; Zhang et al., 2003), are also quantifiable in the periphery. The use of peripheral blood, instead of other tissue types, allows the unique capability of sampling over time points along tumor progression. Such information flow, out of the tumor, and into repertoire data, is taking place through multiple channels. Blood flow in and out of the tumor provides direct tethering between different T cell compartments, and other channels of control over the CDR3 repertoire include interactions between multiple arms of the immune system that interact with the tumor microenvironment (Gajewski, Schreiber, & Fu, 2013).

The co-evolution of the T cell repertoire and a neoplastic tumor over a significant period (9 months) of mice life is accompanied by the evolution of the TCR repertoire during mice maturation and mice ageing. It has already been demonstrated that the T cell repertoire changes significantly with age (Britanova et al., 2014; Yoshida et al., 2017). To control for this factor, we used mice with the same genetic background as the transgenic mice, but are non-transgenic and do not develop tumors.

## Results

We followed the development of spontaneous tumors in a transgenic mouse expressing the inactivated rat neu (Erbb2) oncogene. This mouse model serves as a model of HER2+ human breast cancer (Guy et al., 1992). As a control, we use FVB/NJ strain mice with the same genetic background, but lacking the transgene. We sequenced TCR repertoires in the peripheral bloods of each of the 15 mice drawn at 8 monthly time points (Figure 1). We focused on the largest recoverable subset of T cells, those bearing markers CD4^+^CD62L^+^CD44^−^. The counts of TCRα and TCRβ sequences are presented in Supplementary Table 1. We categorized the following groups: Young (mice 2-5 months old); Old (mice 6 months or older); Control; and Transgenic (Tg): pre-tumor; early-tumor (one month before a palpable tumor); and tumor. Supplementary Table 1 tags samples by these categories.

**Table 1.**
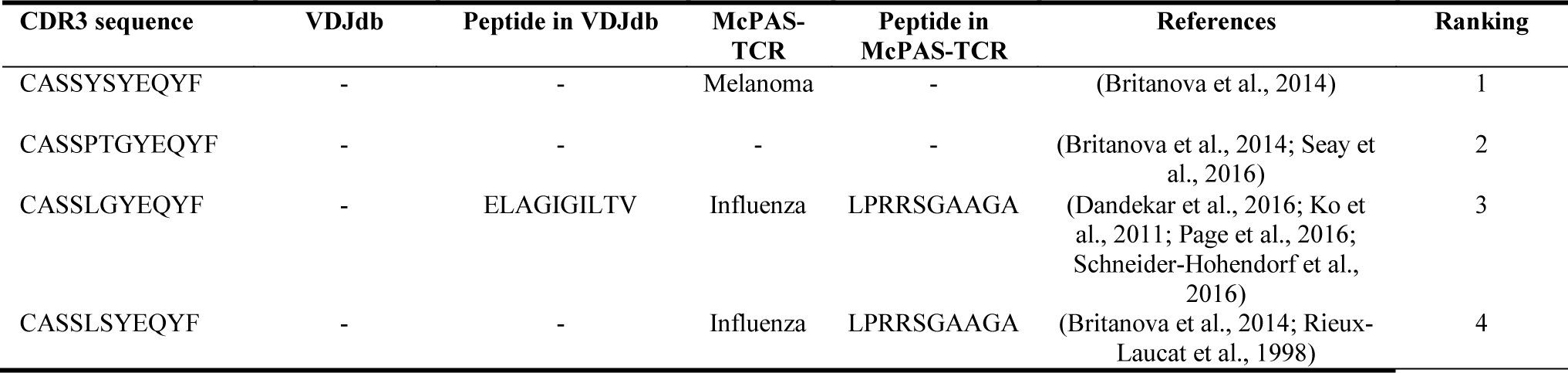
CDR3 sequences that show cross-species, tumor-exclusive, behaviors

**Fig. 1.**
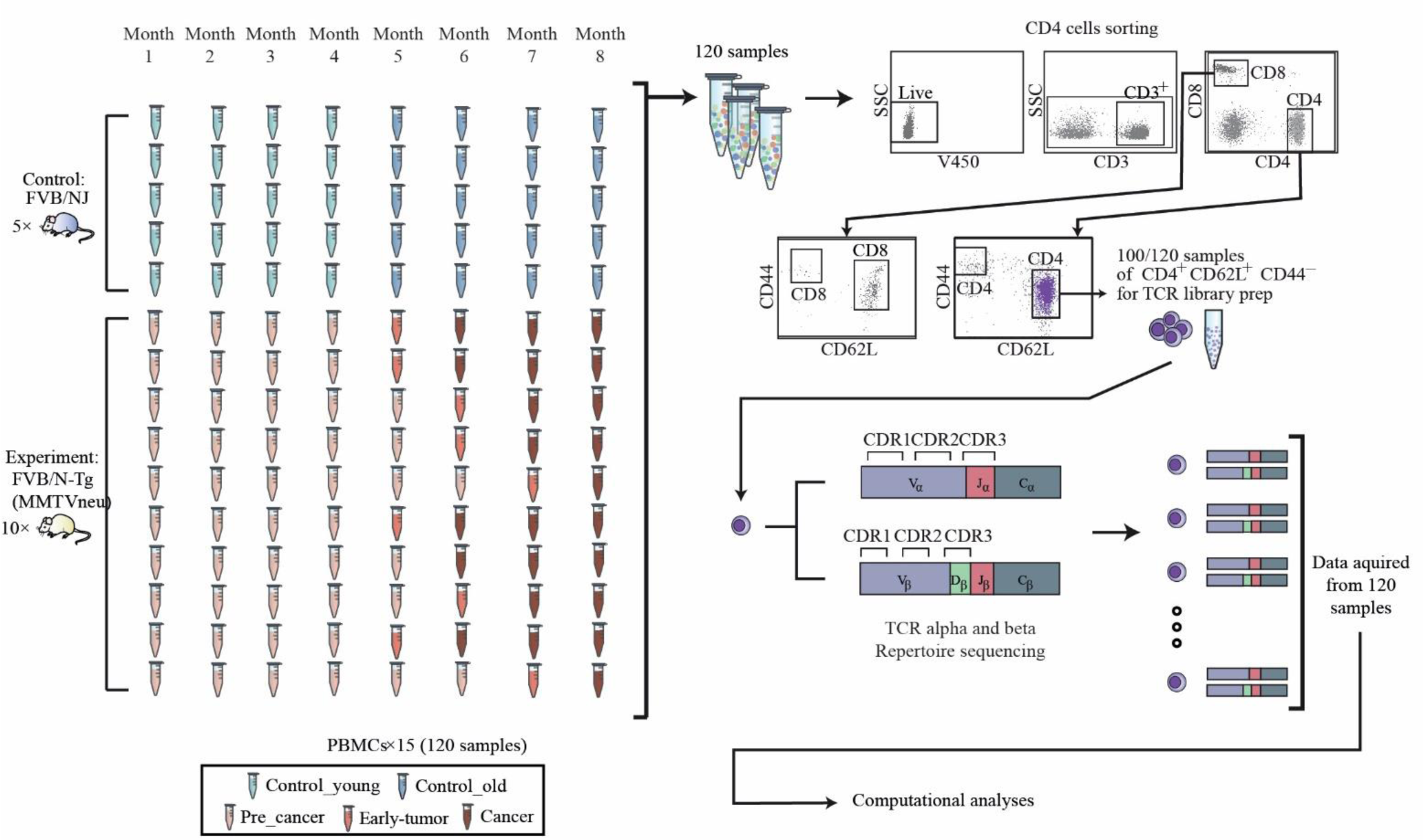
Experimental procedure. 120 blood samples were drawn from the retro-orbital sinus of 10 FVB/N-Tg (MMTVneu), a mouse model of HER2 human breast cancer mice, and from 5 FVB/NJ control mice. Over these 8 time-points, none of the control mice (blue) developed any tumors. Progress of tumor in the ten transgenic mice is demonstrated using the red colored samples in the figure. The last time point before tumors are shown was defined as pre-cancer and marked light red. From each time point, the peripheral blood mononuclear cells were isolated and stained for flow cytometry. Cells were analyzed and gated for sorting using a FACS ARIA III sorter, and CD4+CD62L+CD44- naïve population was separated for RNA extraction and T cell receptor library preparation (see Methods).

We found that there were no significant differences in repertoire diversity or CDR3 length distributions; but there were differences in copy number and percent of unique sequences among the groups (Supplementary Fig. 1). Private clonotypes refer to CDR3 AA sequences appearing in only one mouse; public clonotypes appear in at least two different mice. Public clonotypes are termed *Inclusive* if they appear in more than one phenotype and *Exclusive* if they are shared only by mice of the same phenotype group. We found that Exclusive Public clonotypes are more prevalent in the Tg group (19-23%) compared to the Control group (3-5%; Fig. 2a and Supplementary Fig. 2a). In other words, tumor-exclusive public sequences mark the Tg mice. Interestingly, public clonotypes occupy between 40-60% of the sequences in all the phenotypes.

**Fig. 2.**
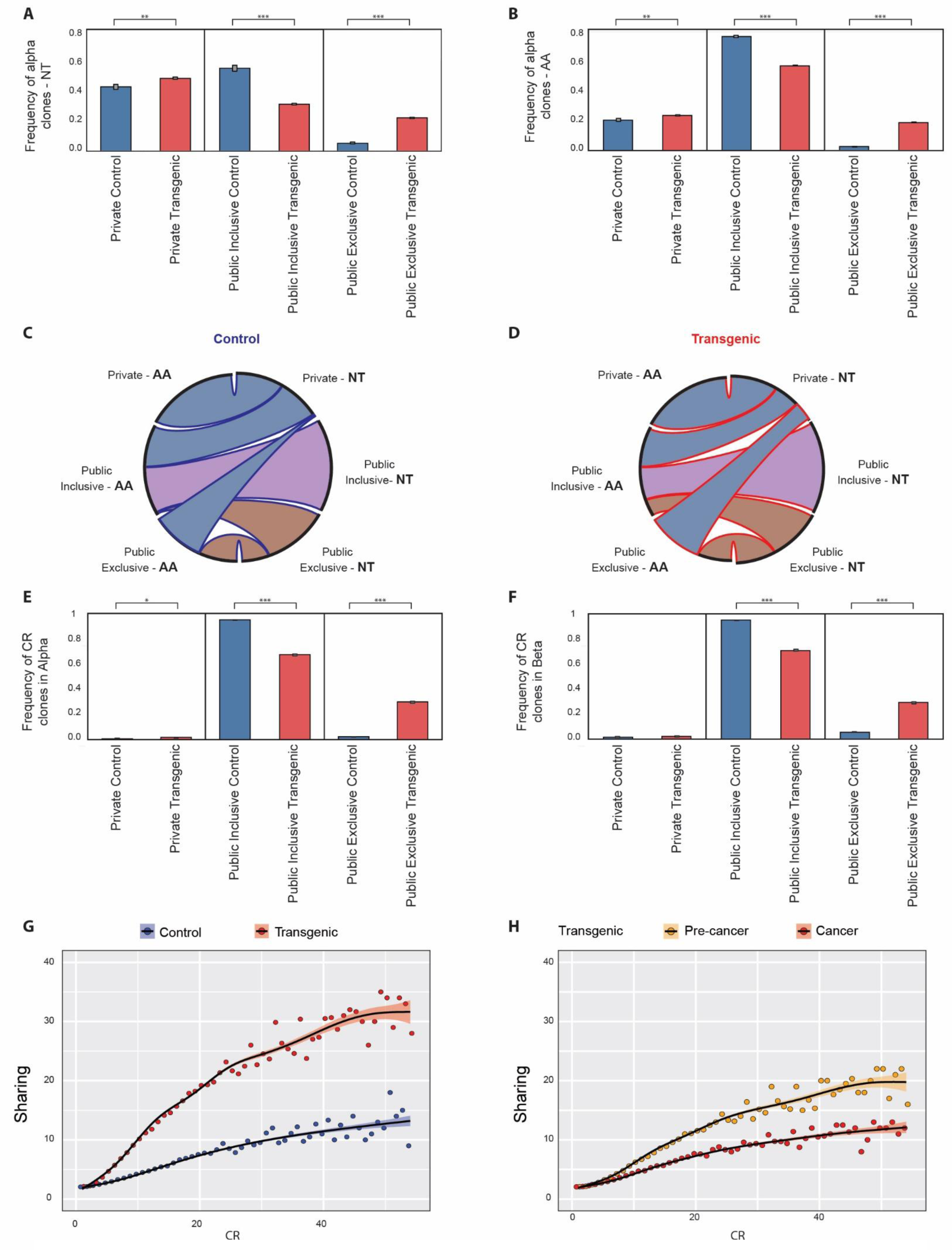
The public repertoire and convergent recombination. Public repertoires and their subtypes are shown in Tg mice and in Control mice using nucleotide sequences (**A**) and using amino-acid sequences (**B**). The two panels show the repertoire in the alpha chain, but a similar effect is seen in the beta chain (Supplementary Fig. 2). **(C, D)** Convergent recombination in control and transgenic mice. The right-hand side of each circle indicates nucleotide (NT) sequences, while the left-hand side indicates amino-acid (AA) sequences. The edges between NT and AA represent the effect we see in convergent recombination – in which different NT sequences encode to the same AA sequence and change the Public/Private balance. **(E, F)** Frequencies of CR clones in the public repertoire in alpha and beta chains. **(G, H)** Correlation between sharing level and CR level in Control versus Tg groups (G) and in No-Cancer group vs. Cancer group (H).

Convergent recombination (CR) refers to identical TCRα or TCRβ amino-acid (AA) sequences originating from different nucleotide (NT) sequences. We found a higher frequency of CR in the α chain (56%- 69%; Supplementary Fig. 3a) compared to the β chain (43%-52%; Supplementary Fig. 3b). Figures 2a and 2b show that CR in the Inclusive Public groups is double that of the Private groups; Figures 2c and 2d show that the Exclusive-Public clonotypes originated from a smaller portion of the private clones in the Control group compared to the Tg group. In the private groups, the non-CR clonotypes dominated the repertoire (Supplementary Fig.3c, 3d). In addition, we identified a significant contribution of CR to the public TCR repertoire – 97%- 99% of the CR clonotypes of the α chain are public (Fig. 2e) compared to 13%-28% of public clonotypes in non-CR clonotypes (Supplementary Fig.3c). The same phenomenon is observed in β chain – 96%-99% of the CR clonotypes are also public clonotypes (Fig. 2f), in comparison to 6%-26% of public clonotypes in the non-CR clonotypes (Supplementary Fig. 3d).

As CR levels increased, the sharing level for all groups increased (Fig. 2g, 2h). Furthermore, the connection between CR and sharing is group specific. In the Tg group, sharing increases faster when CR increases, compared with the Control group (Fig. 2g). This may indicate that selection (associated with CR levels) of specific clonotypes, is dominant in the transgenic mice, perhaps due to the sharing of tumor antigens among mice. Separating the transgenic group into pre-tumor and tumor samples, we see that sharing is greater in the pre-tumor group than in the tumor group (Fig. 2h); thus, high CR and highly shared sequences develop early, before we detect a palpable tumor.

To learn whether our observations in the mouse model might be relevant to human breast cancer, we compared the mouse TCR data to three, previously published human TCR datasets, each of which focused on a different aspect of breast cancer (Figure 3a) (Beausang et al., 2017; Page et al., 2016; Wang et al., 2017): Beausang et al (Beausang et al., 2017) studied T cell receptors in early-stage breast cancer and sampled the tumor tissue, the surrounding normal tissue and peripheral blood. Wang et al (Wang et al., 2017) sequenced the TCR repertoires of breast cancer tumors, lymph nodes and adjacent tissue. Page et al (Page et al., 2016) studied breast cancer after immunotherapy. We asked two questions: do mice and humans with breast cancer share cross-species TCR repertoires, and do these cross-species TCRs manifest common CR events.

**Fig. 3.**
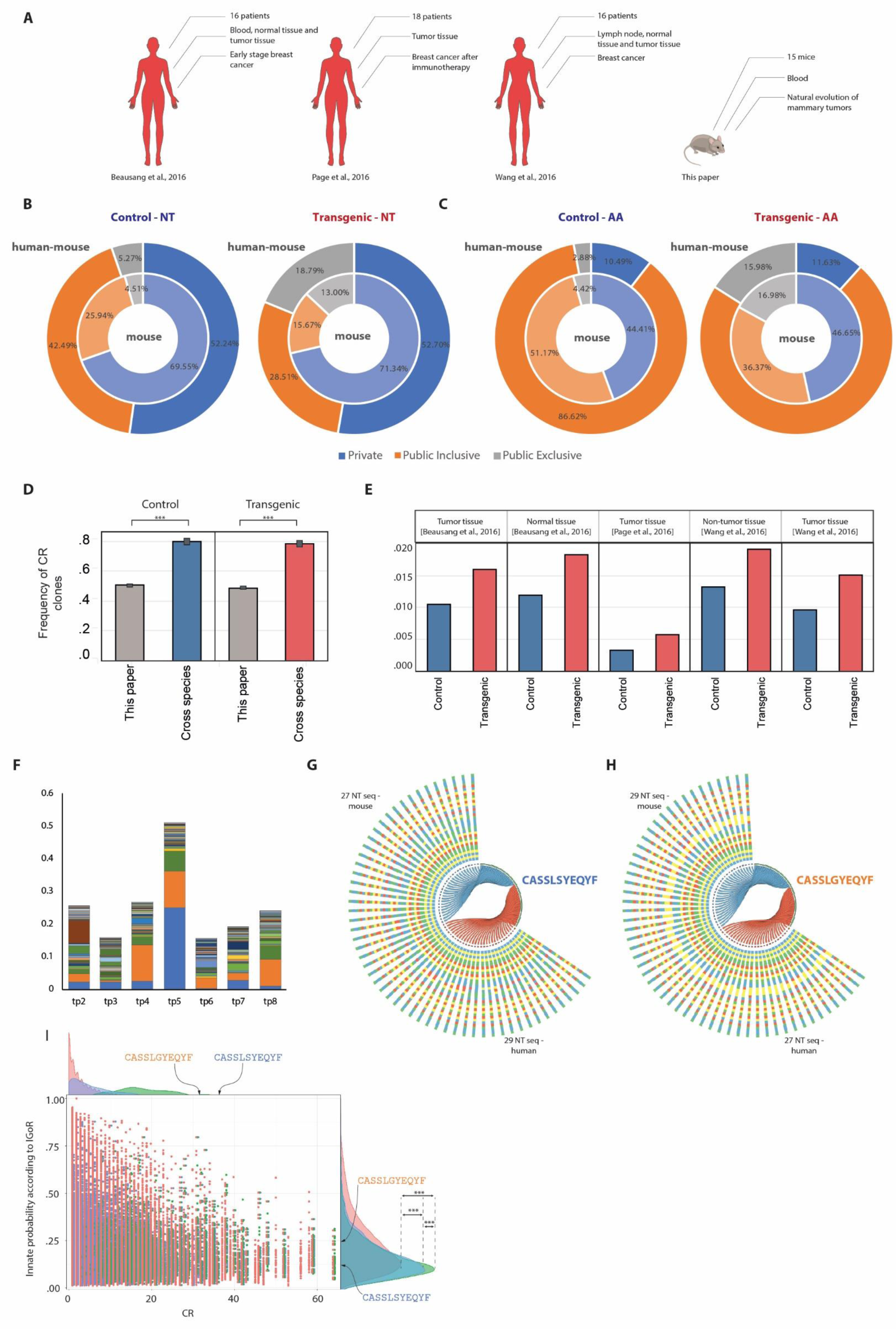
The connection between the mouse repertoire and human breast cancer. To learn more about the connection between the repertoires of mouse mammary tumor and human breast cancer, we studied the three published datasets described in **(A)**. These include 50 breast cancer patients from different studies, in which different conditions and different tissues were studied. (**B, C**) The Public repertoires in the sub-populations of clones shared with human samples are shown using an NT view (**B**) and an AA view (**C**). (**D**) The frequency of CR clones that are shared with the human datasets is higher than the one measured in the mice alone. (**E**) The percentile of clones that are shared between human and mouse samples shows that the Tg share more sequences with human samples (note that all human samples come from tumor-bearing individuals). (**F**) We ranked the 258 cross-species sequences according to their abundance in each sample. To visualize similarities between the ranking in each time point and the samples from early stage breast cancer patients as in Beausang et al., we stacked bars from each of the 258 ranking lines. The area of each bar has been determined so that it is reciprocal to (ranking in human X ranking in mouse). In that manner, if, in a specific time point, a clonotypes is ranked #1 in mouse samples, and is also ranked #1 in human samples, it would demonstrate the largest area. Color of sequences is preserved across bars, so that we see that three sequences dominate the similarity between samples: blue, orange and green. These sequences are included in Table 1, and, as the Table further indicates, have been previously associated with Melanoma, with Influenza, and with Diabetes. **(G, H)** The two highly ranked clones in the cross-species analyses are visualized for their CR sources. In the panel, each nucleotide sequence is connected to its translated amino-acid sequence. Edges are blue if they originate from a mouse sequence and red if they originate from a human sequence. As the panel shows, there is no overlap between sources for these two amino-acid sequences. **(I)** We used IGoR (Marcou, Mora, & Walczak, 2018) (see Text) to learn of any differences between the populations of cross-species sequences used in panels (**F**, **G**, **H**) and the full collection of sequences. To do that, we plotted each sequence as a dot on the graph. Red dots represent cross-species sequences, blue dots represent the full set of all sequences, and green dots represent the 4700 NT sequences that code for the 258 AA sequences that are shared across all time-points and with human samples. The vertical location of the dot is determined by its IGoR value and the horizontal location by its CR value. The right-hand side curves present the histogram over the IGoR values, and the upper-side curves present the histograms over CR values. We used a Kolmogorov–Smirnov test to estimate p-value for the differences between distributions. Indeed, a highly significant p-value (p<2.2×10^−16^) has been obtained, that demonstrates a large difference between the sequence populations. It is interesting to note, as shown by the three CR histograms on the upper side of the panel, that the CR values of these three populations of sequences also come from extremely different distributions (p<2.2×10^−16^). We also highlighted the locations of the two sequences that are described in panels (**F**, **G**, **H**). Although these two sequences differ by a single amino-acid. Their IGoR scores are very different.

As we shall elaborate in greater detail below, mice and humans with breast cancer do share TCR CDR3 AA sequences. Figure 3b compares mouse-specific NT and AA sequences (inner circle) with cross-species mouse sequences shared with human patients (outer circle); the groups are divided into Private, Public Inclusive and Public Exclusive. Regarding NT sequences (the left two wheels), it can be seen that the Tg mice express more Public Exclusive and less Public Inclusive sequences than did the Control mice. AA sequences, in contrast, manifest a much greater proportion of Public Inclusive sequences than Public Exclusive sequences. Nevertheless, Tg mice do express relatively more Public Exclusive sequences. Thus, NT and AA sequences appear to be unconnected generally, and mice developing tumors develop relatively more Public Exclusive sequences of both NT and AA types.

Figure 3d shows that cross species clonotypes shared by mice and humans, both Control and Tg, manifest greater frequencies of CR than do mouse-specific sequences.

Figure 3e, compares the Control and Tg mouse groups to each of the three human studies regarding Tumor, non-tumor and normal tissues. It can be seen that in each case the Tg mouse sequences manifested more sharing with the human repertoires than did the Control sequences. Tumor development appears to amplify the proportion of mouse TCR clonotypes shared with human breast cancer patients, even in non-tumor tissue. Note that the human non-tumor samples were obtained from cancer patients; there was no human Control group equivalent to the mouse Control group.

To obtain an additional metric to match samples across species, we used a ranking system to compare mouse sequences with sequences obtained from early-stage breast cancer patients reported by Beausang et al. First, we filtered for the set of sequences that appear in all mouse time-points and in the human samples. This resulted in 258 CDR3 cross-species sequences. At each time-point, we ranked the sequences according to their copy number– the highest copy number was ranked 1, he second highest ranked 2 and so on. We then asked if the ranking in the human samples was similar to the ranking in the mouse sample, at each time point. To quantify these differences, we produced the bar-chart of Figure 3g. Every time-point in the bar-chart contains a stack of 258 bars. The area each bar occupies in the stack-bar is inversely proportional to an area defined by the ranking of the sequence at the specific time-point, multiplied by the ranking of the sequence in the human samples, such that similarly ranked sequences occupy a larger area. For example, the largest area would be occupied by a sequence ranked 1 in both species. Figure 3f, at time-point 5, which is associated with early-tumors in mice, proves to be the time-point most similar to early-tumors in patients. We can also see that a small set of sequences (blue, orange, green, red in Figure 3f) dominates the chart. When we further study those specific sequences, we see that they are somewhat ubiquitous to TCR repertoire studies. They appear in other works, available from public datasets, and have been associated with Melanoma and Influenza (Table 1). Recent findings from other works show associations to these phenotypes (Laubli et al., 2018; Simoni et al., 2018).

Finally, to learn whether identical CDR3 AA sequences in mice and humans might originate from similar NT recombinations, we selected AA sequences that are highly ranked across species (blue and orange in Figure 3f), and compared their parent NT sequences. The results are shown in Figures 3g and 3h, which connect each NT sequence to its end-result AA sequence. Edges are blue if they originate from a mouse sequence and red if they originate from a human sequence. We can see that the same amino-acid sequence originates from multiple NT sequences, and that the NT sequences have no overlap between the two species. The AA sequence in figure 3h originates from 29 different DNA sequence in the mouse and from 27 different NT sequences in the human; and the AA sequence in Figure 3h originates from 29 different NT sequences in mice and 27 different NT sequences in humans. Whatever the upstream NT recombination, the CR process strongly selects for a specific AA sequence. To learn more about the differences between the specifics groups of cross-species sequences and the general population of CDR3 sequences, we utilized IGoR, which is a recently developed tool (Marcou et al., 2018), that probabilistically annotates sequences. Figure 3i shows the results of IGoR based analyses over the three groups discussed here: the collection of all sequences, the collection of all cross-species sequences, and the collection of the unique set of 258 cross-species sequences that are inherent to all samples, together with human samples. As the figure shows, the three curves are 1) significantly different from each other 2) Cross-species sequences and the set of 258 sequences are associated with a distribution of relatively low probabilities on the one hand, and high CR (displayed at the top of the panel) distributions. These indicate that even though the sequences were not produced at higher probabilities, they were selected to appear as such. Of special interest, and highlighted in the figure, are the two sequences mentioned above, CASSLGYEQYF and CASSLSYEQYF, which, in spite of their single AA difference, appear with very different IGoR scores and with very high CR values.

## Discussion

Investigation of the temporal behavior of TCR repertoires during mammary tumor development in mice, shows that they are dominated by Public clonotypes shared with human breast cancer patients. These shared, cross species TCR sequences are very likely to arise as a consequence of strong selection by antigen epitopes present in both mice and humans with breast tumors, as indicated by the continuous involvement of CR. This CR and public clonotypes shape the unique structure of the repertoire in mice with tumors. The phenomenon of cross-species sequences has been demonstrated before (Madi et al., 2014) and is strongly associated here with Inclusive-public (public only in tumor) sequences. We also showed that early tumor development in mice shares a TCR repertoire with early tumors in human samples and that these specific sequences show high levels of CR, that originate from different nucleotide sequences in mice and different nucleotide sequences in human, with no overlap, further indicating selection related to mammary tumors. Public databases (Shugay et al., 2018; Tickotsky, Sagiv, Prilusky, Shifrut, & Friedman, 2017) show that these specific TCR sequences have previously been seen in connection with specific peptides reported in connection with other cancer types, and with CMV and influenza viruses. The sequences we report on here should be studied further in connection with responses to tumor immunotherapy. Modifying tumor repertoires to shift them in specific directions, which may be quantified according to some of the analytics we show here, may allow for novel treatments.

## Materials and Methods

### Mice

Transgenic Mice expressing the inactivated rat neu (Erbb2) oncogene under the transcriptional control of the mouse mammary tumor virus promoter were purchased from Jackson Laboratories (FVB/N-Tg(MMTVneu) 202 Mul/J). The female mice of this strain represent a mouse model of mammary tumor in humans, model of HER2/ Erbb2 / Neu human breast cancer (Guy et al., 1992). FVB/NJ strain with the same genetic background as the transgenic mice, serve as a non-transgenic control mouse that does not develop tumors. Mice were housed in accordance with all applicable laws and regulations following approval by the responsible animal care and ethical committee, under specific pathogen-free conditions. Mice were monitored by palpitation for tumor development monthly for up to 9 months.

### Antibody staining and cell sorting

Blood was sampled from the retro-orbital sinus of 15 mice once per month for 8 time points (total of 120 samples). Mononuclear cells from the peripheral blood was isolated by density gradient centrifugation using Ficoll (Ficoll PaqueTM plus, GE Health Care), Single cell suspensions were prepared from thymus and spleen that were removed from each mouse at the end of the experiment. For cell sorting, cells were stained with the following fluorescently labeled monoclonal antibodies: anti-CD4 Pacific Blue (BD), anti-CD25 PE (eBioscience), anti-CD44 APC (BD) and anti-CD62L PE-Cy7 (eBioscience) and viability using the Fixable Viability stain 450 (BD Horizon). Cell sorting was performed using FACS ARIA III sorter. CD4+ D44loCD62Lhi were sorted as naïve T cells. After sorting, cells were pelleted and resuspended with 300µl of RNAprotect cell reagent (Qiagen). Cells were stored at minus 80°C until RNA extraction. RNA was purified from RNAprotect-stabilized cells using the RNeasy Plus Mini Kit. After RNA extraction, samples were run on TapeStation to estimate quality.

### High-throughput sequencing of the T cell repertoire

The method for high-throughput sequencing of the T cell repertoire was performed as previously described (Di Niro et al., 2015; Tsioris et al., 2015). Briefly, RNA was reverse-transcribed into cDNA using a biotinylated oligo dT primer. An adaptor sequence was added to the 3′ end of all cDNA, which contains the Illumina P7 universal priming site and a 17-nucleotide unique molecular identifier (UMI). Products were purified using streptavidin-coated magnetic beads followed by a primary PCR reaction using a pool of primers targeting the TCRα and TCRβ regions, as well as a sample-indexed Illumina P7C7 primer. The TCR-specific primers contained tails corresponding to the Illumina P5 sequence. PCR products were then purified using AMPure XP beads. A secondary PCR was performed to add the Illumina C5 clustering sequence to the end of the molecule containing the constant region. The number of secondary PCR cycles was tailored to each sample to avoid entering plateau phase, as judged by a prior quantitative PCR analysis. Final products were purified, quantified with Agilent Tapestation and pooled in equimolar proportions, followed by high-throughput paired-end sequencing on the Illumina MiSeq platform. For sequencing, the Illumina 600 cycle kit was used with the modifications that 325 cycles was used for read 1, 6 cycles for the index reads, 300 cycles for read 2 and a 20% PhiX spike-in to increase sequence diversity.

### Processing of raw sequencing data

Transforming raw reads to alpha and beta CDR3 sequences is done by using MiXCR, a universal framework that processes big immunome data from raw sequences and output quantitated clonotypes (Bolotin et al., 2015). MiXCR performs paired-end read merging and extracts human or animal BCR and TCR clonotypes providing corrections of erroneous sequences introduced by NGS (Kivioja et al., 2012; Vander Heiden et al., 2014; Yaari & Kleinstein, 2015).

